# Sensory stimulus evoked responses in layer 2/3 pyramidal neurons of the hind paw-related mouse primary somatosensory cortex

**DOI:** 10.1101/2020.08.30.274308

**Authors:** Guillaume Bony, Arjun A Bhaskaran, Katy Le Corf, Andreas Frick

## Abstract

The mouse primary somatosensory cortex (S1) processes tactile sensory information and is the largest neocortex area emphasizing the importance of this sensory modality for rodent behavior. Most of our knowledge regarding information processing in S1 stems from studies of the whisker-related barrel cortex (S1–BC), yet the processing of tactile inputs from the hind-paws is poorly understood. We used *in vivo* whole-cell patch-clamp recordings from layer (L) 2/3 pyramidal neurons (PNs) of the S1 hind-paw (S1-HP) region of anaesthetized wild type (WT) mice to investigate their evoked sub- and supra-threshold activity, intrinsic properties, and spontaneous activity. Approximately 45% of these L2/3 PNs responded to brief contralateral HP stimulation in a subthreshold manner, ~5% fired action potentials, and ~50% of L2/3 PNs did not respond at all. The evoked subthreshold responses had long onset- (~23 ms) and peak-latencies (~61 ms). The majority (86%) of these L2/3 PNs responded to prolonged (stance-like) HP stimulation with both on- and off-responses. HP stimulation responsive L2/3 PNs had a greater intrinsic excitability compared to non-responsive ones, possibly reflecting differences in their physiological role. Similar to S1-BC, L2/3 PNs displayed up- and down-states, and low spontaneous firing rates (~0.1 Hz). Our findings support a sparse coding scheme of operation for S1–HP L2/3 PNs and highlight both differences and similarities with L2/3 PNs from other somatosensory cortex areas.

**KEY POINTS:**

- Responses of layer (L) 2/3 pyramidal neurons (PNs) of the primary somatosensory hind-paw cortex (S1-HP) to contralateral hind-paw stimulation reveal both differences and similarities compared to those of somatosensory neurons responding to other tactile (e.g. whiskers, forepaw, tongue) modalities.
- Similar to whisker-related barrel cortex (S1-BC) and forepaw cortex (S1-FP) S1-HP L2/3 PNs show a low spontaneous firing rate and a sparse action potential coding of evoked activity.
- In contrast to S1-BC, brief hind-paw stimulus evoked responses display a long latency in S1-HP neurons consistent with their different functional role.
- The great majority of L 2/3 PNs respond to prolonged hind-paw stimulation with both on- and off-responses.
- These results help us to better understand sensory information processing within layer 2/3 of the neocortex and the regional differences related to various tactile modalities.

## INTRODUCTION

The sense of touch facilitates exploration, recognition of the environment, texture discrimination, sensory-motor feedback, and social interaction (Abraira and Ginty, 2013). In rodents, the primary somatosensory cortex (S1) processing this tactile sensory information is — with a surface area corresponding to ~25% of the entire neocortex — the largest sensory neocortical area, highlighting the importance of this sensory modality for rodent behavior (Catania and Remple, 2002; Seelke et al., 2012). Tactile signals are generated by activation of various types of touch receptors (Roudaut et al., 2012; Zimmerman et al., 2014) and reach the neocortex mainly through the dorsal column pathway, the brainstem, and then via the thalamus (Abraira and Ginty, 2013). The processing of tactile sensory information in the neocortex has been extensively studied for the whisker system and the related barrel cortex (S1-BC; reviewed in (Brecht, 2007; Feldmeyer, 2012; Petersen and Crochet, 2013). Recent studies have described the peripheral sensory neurons of fore- and hind paws mediating various aspects of innocuous tactile information (Abraira and Ginty, 2013; Bai et al., 2015; Severson et al., 2017; Walcher et al., 2018). However, knowledge of the neocortical processing of tactile information derived from the HPs, however, is largely lacking. In rodents, the HPs are involved in several behaviors, such as grooming, postural reflexes, walking, and swimming (Whishaw et al., 1999).

Within layer (L) 2/3 of S1, sensory information spreads both vertically within the home cortical column and horizontally across neighboring columns (reviewed in (Feldmeyer, 2012). In addition, these layers receive information from, and project to, many different neocortical regions. Thus, L2/3 PNs of S1-HP are key integrators of information across cortical regions. L2/3 PNs of S1–BC are characterized by a sparse action potential (AP) firing code even in response to a given whisker stimulus (e.g. (Barth and Poulet, 2012; Brecht et al., 2003; Crochet et al., 2011; de Kock et al., 2007; de Kock and Sakmann, 2009; Margrie et al., 2002). Here, we explored whether sensory processing features in L2/3 PNs of S1-HP such as supra- and sub-threshold responses, the latencies, and on-/off-components of the HP related information are unique for this tactile modality or more broadly applicable across different tactile modalities. For example, a sparse firing rate has been suggested to be a general landmark of L2/3 PNs of primary sensory cortices (Barth and Poulet, 2012), but this has never been shown for S1-HP. Further, we asked if neurons that respond to sensory stimulus and those that do not could be distinguished based on their intrinsic properties. We addressed these questions by performing whole-cell patch-clamp recordings together with HP stimulation in anesthetized mice.

## METHODS

### Ethical approval

All experimental procedures were performed in accordance with the EU directive 2010/63/EU and French law following procedures approved by the Bordeaux Ethics Committee (CE2A50). WT mice from our transgenic breeding program (as described in (Zhang et al., 2014) were used in these experiments. These mice were generated by backcrossing 129/Sv/C57Bl/6J/FVB founders into a C57Bl/6J background (6 generations). These mice were chosen to permit comparison with transgenic mice for a companion study and to spare the number of mice used in the two studies.

### Animal preparation and hind paw stimulation

Male WT mice (P24–32) were anaesthetized with a mixture of ketamine (100 mg.kg^-1^) and xylazine (10 mg.kg^-1^) injected intraperitoneally. Mice were monitored for whisker movements, eye blinking, tail and toe pinch reflexes and anesthesia was supplemented if necessary, throughout the experiment. Mice were head-fixed using non-puncture ear-bars and a nose-clamp (SR-6M, Narishige). Body temperature was maintained at 37°C. A small craniotomy was made above the S1-HP (1 mm posterior, 1.5 mm lateral from Bregma) using a dental drill (World Precision Instruments). The stereotaxic coordinates were assessed in a set of control experiments using flavoprotein autofluorescence imaging.

Sensory responses were evoked by applying current pulses (2 ms or 200 ms, 100 V, 0.5–30 mA) via conductive adhesive strips (~1 cm^2^) placed on top of, and below the HP, as described previously (Palmer et al., 2012). These electrodes cover the entire paw (digits and palm, both glabrous and hairy skin). We repeated the stimulation protocol 40 times at a rate of 0.3 Hz.

### *In vivo* whole-cell patch-clamp recordings

Blind whole-cell patch-clamp recordings were performed from L2/3 PNs as described previously (Margrie et al., 2002). Pipettes with an open-tip resistance of 4–6 MΩ were pulled from borosilicate glass using a PC-10 puller (Narishige) and filled with internal solution composed of (in mM): 130 K-methanesulfonate, 10 Hepes, 7 KCl, 0.05 EGTA, 2 Na2ATP, 2 MgATP, 0.5 Na2GTP; pH 7.28 (adjusted with KOH). In a subset of experiments, biocytin (3 mg/ml) as described in (Rancz et al., 2011) was added to the recording solution for *post-hoc* neuronal identification. The intracellular solution was filtered using a 0.22-μm pore-size centrifuge filter (Costar Spin-X). Signals were acquired using a Multiclamp 700B amplifier and Clampex 10.4 software (Axon Instruments). Data were low pass filtered at 3 kHz and sampled at 20 kHz.

### Staining of neuronal morphology and neocortical barrels

Following biocytin filling of the recorded neurons, the brains were fixed by transcardial perfusion with 4% paraformaldehyde in PBS, and post-fixed for 2h in the same solution. Subsequently, 80-μm-thick tangential slices were cut using a vibratome (Leica), and biocytin was detected using streptavidin-Alexa Fluor 555 (1:1,000, 2h at room temperature (RT)). To visualize the S1 barrels, we performed immunostaining against GAD67 in 3 mice (Meyer et al., 2011) using the following protocol: Mouse on mouse (MOM) blocking/permeabilization (0.3% triton X-100, 4% NGS, and 3% BSA in PBS) for 30 min at RT on a shaker; primary antibody (Mouse anti-GAD67 clone 1G10.2; 1:1,500; Millipore) overnight at 4°C; secondary antibody (Goat anti-Mouse IgG2A Alexa 488; 1: 500; Life technologies) for 2h at RT. Images were acquired with a confocal microscope (Leica) or a slide scanner (Nanozoomer, Hamamatsu), and S1 reconstructions were performed with ImageJ software (NIH).

### Data analysis

To measure input resistance, we injected 500-ms-long hyperpolarizing (−100 pA) square current pulses and measured the membrane potential deflection at 300 ms relative to baseline. AP threshold was measured for the first AP occurring during an IV curve protocol consisting of a series of 500-ms-long current injections ranging from −400 pA to +550 pA (step size: 50 pA). AP half-width was determined by measuring the duration of the first AP at half-amplitude (from threshold to the peak) (see *Figure 1D*). We determined average up- and down-state membrane potentials by plotting the distribution of membrane potential values. Up-state frequency and duration analysis were adapted from (Beltramo et al., 2013). Briefly, the spontaneous down- to up-state transitions were identified as membrane depolarization crossing a threshold set at 1/3 of the amplitude down- to up-state. Only transitions in which the signal remained for more than 150 ms above the threshold were considered. Parameters of evoked sub-threshold synaptic potentials were calculated from an averaged trace of 40 successive trials. Latency was determined by measuring the time point after HP stimulation where the Gaussian fit of the response’s rising phase crosses the baseline. The duration of the synaptic responses was calculated by measuring the width of the averaged response at half-maximal amplitude. The calculation of evoked supra-threshold responses and their coefficient of variation were adapted and modified from (de Kock et al., 2007). Briefly, the number of APs was quantified within a 200-ms-long time window following the stimulus onset and averaged over 40 stimulus trials.

**Figure 1.**
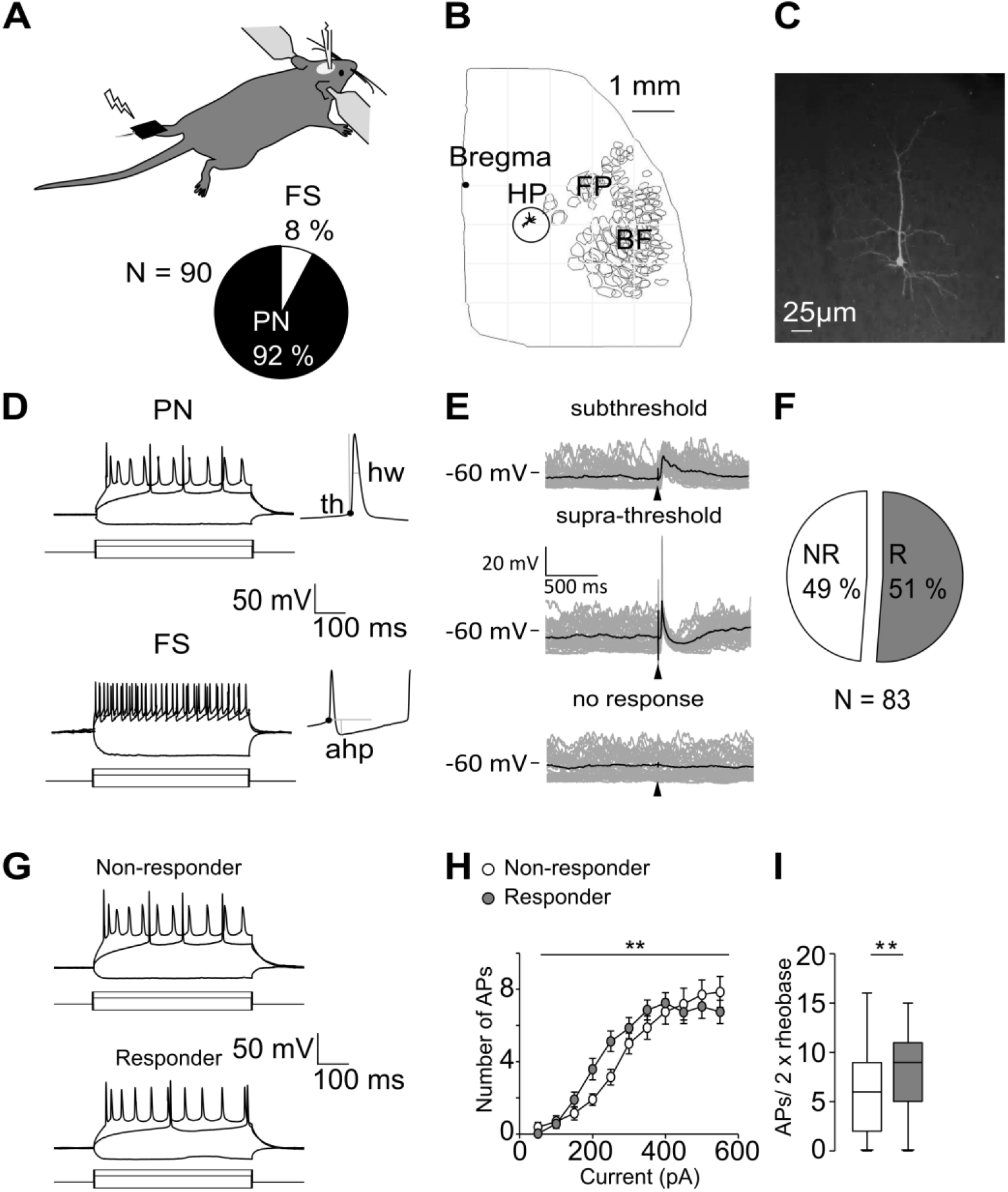
Physiological properties of L2/3 pyramidal neurons in S1-HP of anaesthetized mice. (A) Experimental design: Whole-cell recordings were made from L2/3 neurons of S1-HP while stimulating the contra-lateral HP (top). Proportion of different neuron types recorded (bottom); PN: pyramidal neurons; FS: Fast-spiking interneurons. (B) Position of a recorded and biocytin-filled L2/3 PN within S1-HP. This neuron did not respond to HP stimulation. The circle shows the HP region based on the Allen Brain atlas. HP, hind paw; FP, forepaw; BF, barrel field. (C) Morphology of the same neuron. (D) Example traces from an I-V curve of a pyramidal neuron and a fast-spiking interneuron. Illustration of AP threshold (th), AP half-width (hw) and afterhyperpolarization (ahp) of the first AP for each neuron (inset). (E) Example traces of subthreshold responses over 40 trials (light color) superimposed with the average response (dark color) in an individual L2/3 PN (top); example of suprathreshold responses over 40 trials (middle); example of a non-responder cell over 40 trials (bottom). The arrowhead indicates the HP stimulation (2 ms, 30 mA). (F) Fraction of L2/3 PNs responding to HP stimulation (responding neurons (R), n = 42; non-responding neurons (NR), n = 41). (G) Example traces from an I-V curve of an R- and an NR-cell (−200 pA, rheobase, rheobase x 2). (H) Average number of APs as a function of current injected for R- and NR-cells. (I) Average number of APs at two times rheobase for R- (n = 35) and NR-cells (n = 30). Box plots show the median, interquartile, and range. Statistical significance was calculated by two-way ANOVA with repetition or unpaired Student’s t test. **P < 0.01 (R-cells compared to NR-cells).

The average spontaneous activity (0–200 ms window before stimulus) was then subtracted from this value. The coefficient of variation was calculated by dividing the number of APs within 200 ms following the stimulus by the standard deviation on a trial-by-trial basis. To measure on- and off-responses we used a 200-ms-long stimulation (30 mA, 100 V). All analysis was performed using Clampfit software (Axon Instruments).

### Statistical analysis

Data are given as means ± SD unless otherwise stated. Statistical analysis was performed using two-tailed unpaired or paired *t* tests to evaluate the difference between two groups of data. For repeated measures, data were analyzed by two-way ANOVA (GraphPad Software). P values < 0.05 were considered significant (* P < 0.05, ** P < 0.01, *** P < 0.001). Boxplots indicate the median value (middle line), the 25^th^ and 75^th^ percentiles (box), and the highest and lowest values (whiskers).

## RESULTS

We recorded the integrative properties, spontaneous activity, and sensory stimulus evoked responses in L2/3 PNs (average sub-pial depth 207 ± 60 μm, n = 83 neurons from 68 mice) of the hind paw (HP) somatosensory cortex (S1-HP) of ketamine-xylazine anesthetized mice using whole-cell patch-clamp recordings *in vivo* (*Figure 1*). Pyramidal neurons were filled with biocytin for *post-hoc* identification (*Figure 1C*). Occasional (~8%; *Figure 1A, D*) recordings from fast-spiking interneurons could be readily distinguished from those of pyramidal neurons based on the neurons’ action potential (AP) properties (Table 1) and were excluded from the analysis.

**Table 1.**

|  | **Pyramidal neurons**<br>**(n = 83)** | **Fast-spiking interneurons**<br>**(n = 7)** | *P-value* |
| --- | --- | --- | --- |
| **RMP (mV)** | -71.9 ± 7.9 | -68.3 ± 12.7 | *P* = 0.2796 |
| **Rin (MΩ)** | 91.3 ± 54.3 | 134.2 ± 53.5 | *P* = **0.0495\*** |
| **APth (mV)** | -30.6 ± 9.5 | -34.7 ± 12.8 | *P* = 0.2972 |
| **APhw (ms)** | 2.1 ± 0.5 | 1.2 ± 0.3 | *P* < **0.0001 \*\*\*** |
| **AHP (mV)** | 3.9 ± 2 | 12.8 ± 4.6 | *P* < **0.0001 \*\*\*** |
| **Rheobase (pA)** | 203.7 ± 108.2 | 142.9 ± 93.2 | *P* = 0.1536 |
| **Max firing (Hz)** | 19.2 ± 7.5 | 59.4 ± 30.8 | *P* < **0.0001 \*\*\*** |

### HP stimulation evoked responses in ~50% of L2/3 pyramidal neurons

Brief stimulation of the contralateral HP (2-ms duration, 30 mA) evoked sub- and supra-threshold responses in ~51% of the L2/3 PNs (42/83 neurons), while ~49% of the neurons (41/83 neurons) failed to respond (*Figure 1E, F*). Several possibilities might explain the lack of response in approximately half of the recorded neuronal population. To address this, we first verified that non-responding neurons were not differentially distributed within L2/3 when compared to responding neurons (distance from pia; responding cells: 228 ± 65 μm; non-responding cells: 228 ± 73 μm; *p* > 0.05). Second, both neuron types were located within S1-HP (for an example of a non-responding neuron in S1-HP, see *Figure 1B, C*). Another explanation for a lack of response could be that the stimulation intensity was below the threshold for evoking a response in these neurons. To examine this possibility, we measured the minimum stimulus intensity required to elicit a response in responding neurons. On average this threshold intensity was ~11 mA (11.1 ± 9.8 mA, n =18) and far below the intensity (i.e. 30 mA) used for our experiments. However, we are not ruling out the possibility that some non-responding cells have a threshold higher than 30 mA.

We then probed whether we could distinguish these morphologically (dendritic branching, data not shown, n = 7 neurons each) similar L2/3 PNs sub-populations based on their integrative properties, spontaneous firing rates, and up- and down-states. Analysis of the passive and active membrane properties revealed that most parameters were comparable (Table 2; *Figure 1G-I*, Supple Figure 1). These include the resting membrane potential (RMP) during down-states as well as during up-states, the input resistance (RN) in down-states, the lack of a sag response (Supple Figure 1), the properties of single APs such as threshold, half-width (Supple Figure 1*D-F*) and rheobase (data not shown). In contrast, the number of APs triggered as function of current injected was larger in responding- as compared to non-responding cells (*P* = 0.0172) (*Figure 1G-H*). Accordingly, at two times rheobase the number of APs was approximately 40% higher for responding cells (R-cells: 8.1 ± 4.0; NR-cells: 5.7 ± 4.0; *P* = 0.0175; n = 35 and 30, respectively) (*Figure 1I*). These findings suggest that the responding cell sub-population might be more excitable compared to the non-responding one.

**Table 2.**

|  | **Subthreshold**<br>**Responders (n = 38)** | **Non-responders (n = 41)** | *P-value* |
| --- | --- | --- | --- |
| **RMP (mV) downstate** | -72.3 ± 8.4 | -71.7 ± 7.5 | *P* = 0.74 |
| **RMP (mV) upstate** | -63.0 ± 11.3 | -61.9 ± 10.2 | *P* = 0.69 |
| **Upstate duration (ms)** | 589.0 ± 208.2 | 488.2 ± 175.4 | *P* = 0.08 |
| **Upstate frequency (Hz)** | 0.9 ± 0.3 | 1.0 ± 0.2 | *P* = 0.12 |
| **Spontaneous firing rate (Hz)** | 0.1 ± 0.2 | 0.2 ± 0.2 | *P* = 0.25 |
| **Rin (MΩ) downstate** | 99.8 ± 48.8 | 89.5 ± 65.4 | *P* = 0.44 |
| **AP threshold (mV)** | -28.6 ± 11.0 | -27.6 ± 11.7 | *P* = 0.71 |
| **AP halfwidth (ms)** | 2.1 ± 0.5 | 2.2 ± 0.7 | *P* = 0.58 |
| **Rheobase (pA)** | 195.1 ± 89.3 | 211.9 ± 123.4 | *P* = 0.48 |
| **AP firing at 2x rheobase** | 8.1 ± 4.0 | 5.7 ± 4.0 | *P* < 0.05 * |

Somewhat surprisingly, this increased excitability is not correlated with any differences in spontaneous firing rate, or up-/down state properties (Table 2, Supple Figure 1 *A-C*). Taken together, the lack of contralateral hind paw stimulus evoked responses together with a lower intrinsic excitability suggests that these L2/3 PNs that might require other forms of tactile stimuli or sub-serve other physiological roles related to perception of temperature or pain (Cain et al., 2001; Milenkovic et al., 2014; Paricio-Montesinos et al., 2020; Walcher et al., 2018). Since the goal of our study was to describe the physiological properties of L2/3 PNs responding to paw stimulus, we did not further investigate these possibilities. The remainder of the manuscript describes the properties of neurons that responded to HP stimulus.

### Spontaneous firing activity and up-/down-states of HP stimulus responding neurons

During quiet wakefulness and under anesthesia, the membrane potential of neocortical pyramidal neurons in the whisker related barrel cortex displays low-frequency fluctuations (up- and down states (Petersen et al., 2003; reviewed in (Castro-Alamancos, 2009; Poulet and Crochet, 2018). Up-states represent brief episodes of increased neocortical activity resulting in strong depolarization of the neurons. In our recordings, up-states had on average a duration of 589 ± 208 ms (n = 35) and occurred with a frequency of 0.9 ± 0.3 Hz (n = 35) (Supple Figure 1*C*), consistent with findings from a recent study in rats (Palmer et al., 2014). During some up states, neocortical neurons fire spontaneously (i.e. in the absence of external stimuli) APs at rates that are cell-type specific (Barth and Poulet, 2012). We found that for L2/3 pyramids of S1-HP, only ~40% spontaneously fired APs (15/38 of cells), and the spontaneous firing rate was very low (Supple Figure *1A, B*) (0.1 ± 0.2 Hz).

### HP stimulus evoked subthreshold responses

Next, we characterized the contralateral HP stimulus evoked sensory responses by triggering 40 successive trials using the maximum stimulus strength as described above (i.e. 30 mA; *Figure 2*). 90% of the neurons responded to contralateral HP stimulation in a subthreshold manner with occasional failures, and only ~10% (4 out of 42) of the neurons responded with an AP in some of the trials (see below). *Figure 2A* shows a representative example of a L2/3 PNs neuron responding to HP stimulations in a subthreshold manner. At the population level, the average amplitude of the subthreshold excitatory postsynaptic potential (EPSP) was 9.0 ± 5.1 mV (median 7.7 mV), ranging from 2.5 mV to 21.7 mV (n = 33) (*Figure 2C*). The amplitude of the EPSP was correlated with the membrane potential prior to the response (*Figure 2D*) and was significantly smaller during up-states as compared to down-states (Regression analysis, R square = 0.3499, n = 38, *P* < 0.001). The slope of rise (measured at 20-80% of the amplitude) of the EPSP was 0.6 ± 0.5 mV.ms (n = 36), the half-amplitude duration was 98.2 ± 76.2 ms (n = 36) (*Figure 2E*), and the area was 833.1 ± 560.4 mV.ms (n = 30).

**Figure 2.**
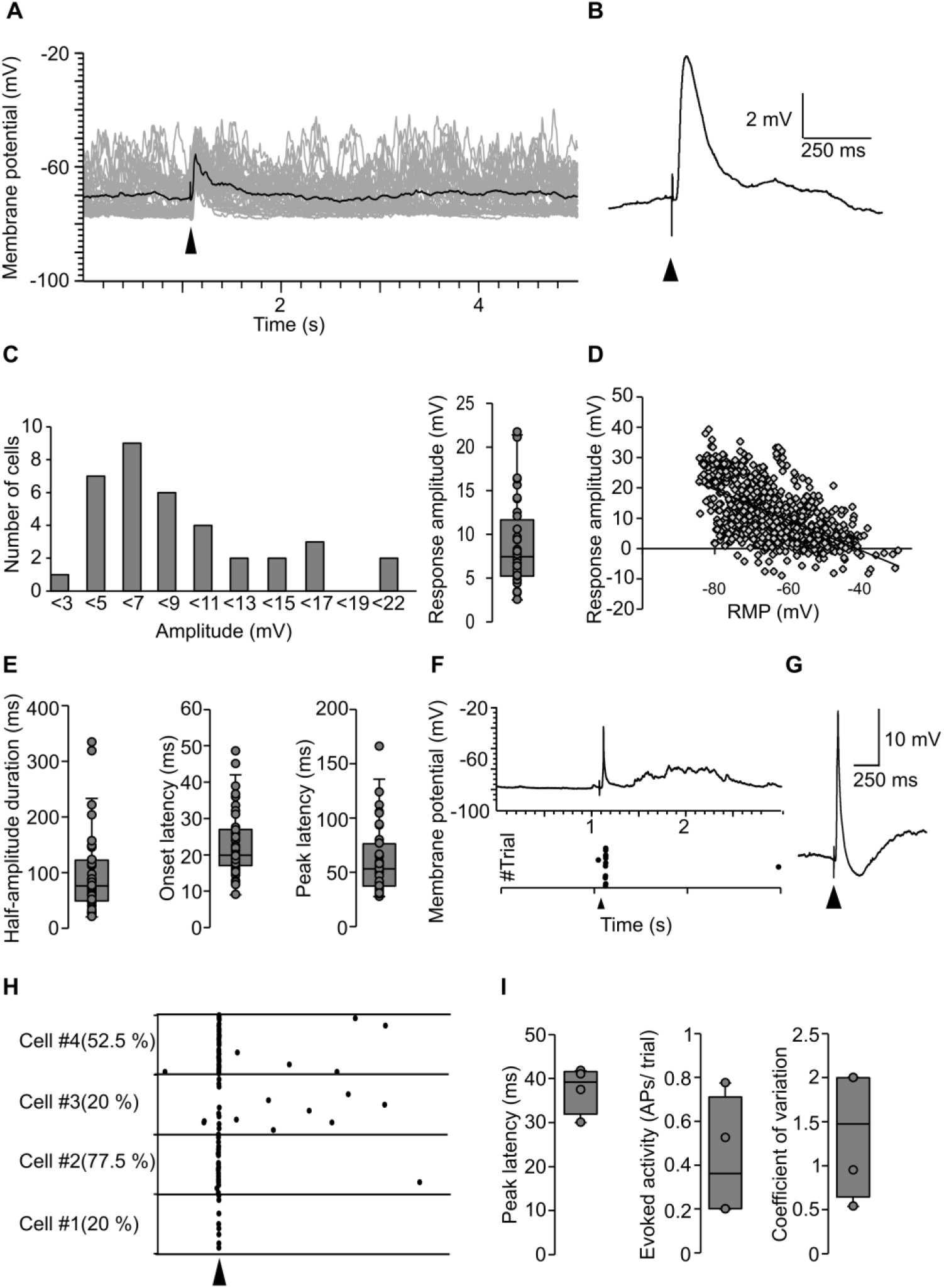
Sub- and supra-threshold responses to HP stimulation. (A) Example traces of subthreshold responses over 40 trials (grey traces) superimposed with the average response (black trace) in an individual L2/3 PN. The arrowhead indicates the time of HP stimulation (2 ms, 30 mA). (B) Grand average of the subthreshold responses of all recorded neurons (n = 36 neurons, 1440 trials). (C) Average amplitudes of subthreshold responses for the whole neuronal population in rank (left) and distribution (right). (D) Correlation between EPSP amplitudes and membrane potential (P < 0.001). (E) Distribution of half-amplitude duration (left), onset latency (middle) and peak latency (right) of EPSPs (n = 36). The latencies were measured from the beginning of the stimulation. (F) Example of suprathreshold recordings (single trial, top); and action potential timing over 40 trials (bottom). Shown are only the traces were APs were triggered. The arrowhead indicates the stimulation onset. (G) Grand average of HP stimulus evoked responses for neurons that responded in a suprathreshold manner in some of the trials (4 cells, 160 trials). (H) Timing of action potentials with respect to HP stimulation. The numbers in parenthesis indicate the percentage of trials with action potential response. (I) Distribution of peak latency (left), evoked activity (middle), and coefficient of variation (right). Box plots show the median, interquartile, range, and individual values.

The onset latency of the EPSP was 21.9 ± 8.9 ms, and the peak latency 58.8 ± 30.8 ms (n = 36) (*Figure 2E*).

### HP stimulus evoked supra-threshold responses in small fraction of neurons

As mentioned above, HP stimulation triggered APs (supra-threshold responses) only in a small fraction of neurons (4/42 neurons responding to HP stimulus). *Figure 2F* shows 40 successive trials of a supra-threshold responding neuron. On average, these neurons responded with a single AP to HP stimulation in 42.5 ± 27.9 % of the trials (range: 20.0% −77.5%) and with a sub-threshold depolarization in the remaining trials (*Figure 2F-I*). Interestingly, HP stimulus evoked APs were followed by an inhibitory component (grand average in *Figure 2G*; 4 cells, 160 trials), as a result of feedforward and feedback inhibition (Isaacson and Scanziani, 2011; Petersen, 2019; Poulet and Crochet, 2018).

### On- and off responses of stance like stimuli

In rodents, the paws are mostly used for sensory/discriminative aspects and for locomotive behavior involving their synchronized movement (Whishaw et al., 1999). Locomotion can be divided into different phases — stance (when the paw is in contact with the floor) and swing (when the paw is moving forward to a new position), and the timing of each phase can vary depending on the animal’s speed. In mice, the average stance phase duration is ~200 ms (Clarke and Still, 1999). To mimic the stimulation received during the stance phase of locomotion, we applied a 200 ms-long stimulus to the HP. The majority of L2/3 PNs responded to both the onset and offset of the stimulus (on- and off responses: 86%; on-only responses: 14% of neurons; n = 14) (*Figure 3A*), with the membrane potential remaining slightly depolarized during the time course of the stimulation (*Figure 3B, C*). The presence of onset and offset responses could reflect either the activation of fast adapting receptors in the skin or it could be a property of neocortical circuits (see Discussion). To describe sensory integration during stance and swing phases, we compared the on- and off-responses (*Figure 3D*). On- and off-responses had similar amplitude (on: 7.9 ± 5.3 mV, n = 7; off: 6.9 ± 3.2 mV, n = 7). The onset latency, however, was significantly longer for off-compared to on-responses (on: 22.1 ± 10.4 ms, n = 7; off: 27.5 ± 9.2 ms, n = 7; *P* = 0.0242), which highlights different processing for these two sensory signals. Peak latency (on: 50.6 ± 21.4 ms, n = 7; off: 62.4 ± 15.5 ms, n = 7) was similar, but half-amplitude duration tended to be longer for off-responses (on: 62.9 ± 15.0 ms, n = 7; off: 89.4 ± 36.6 ms, n = 7; *P* = 0.08) (*Figure 3D*).

**Figure 3.**
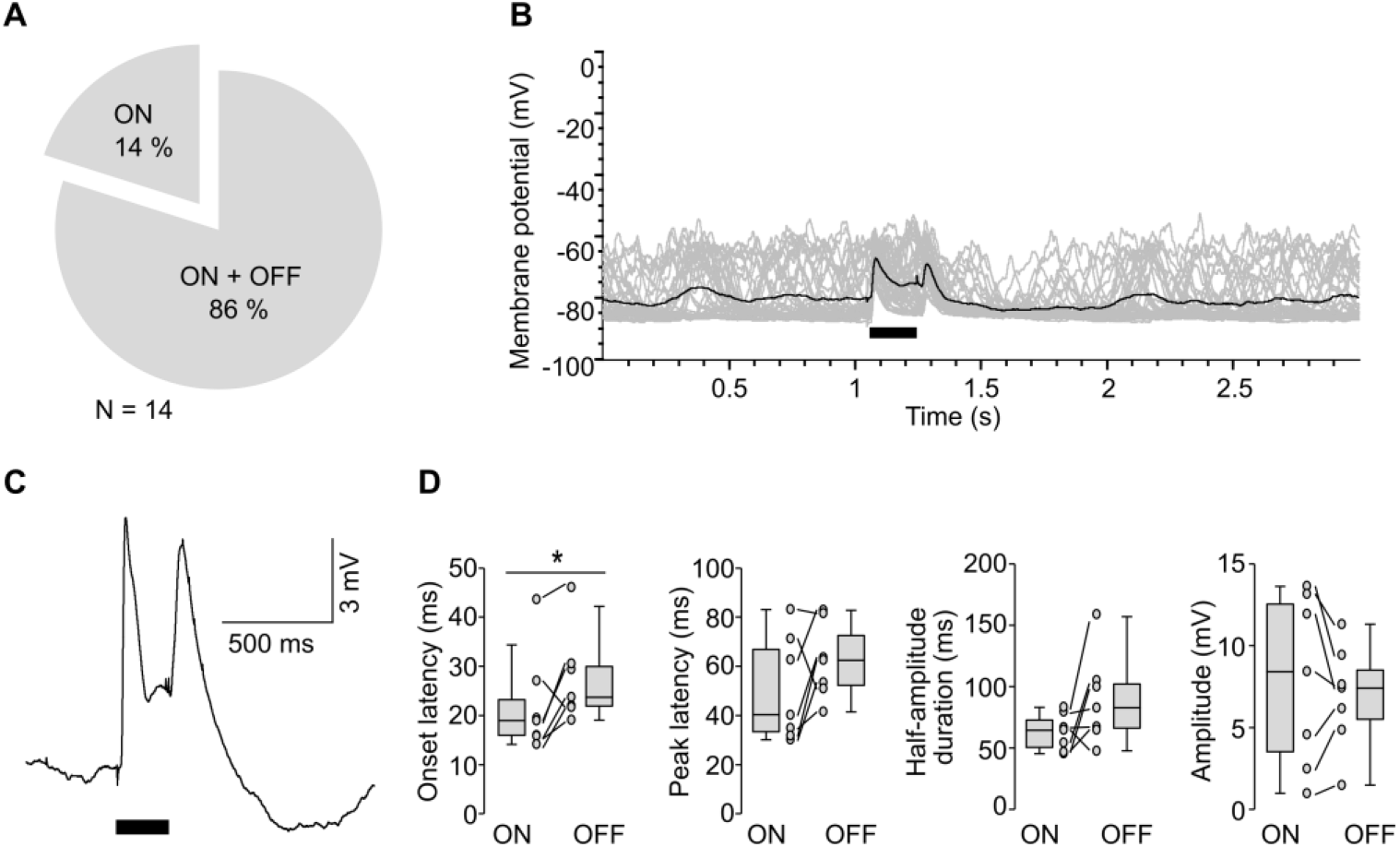
L2/3 pyramidal neurons respond with on- and off-responses to long HP stimulus. (A) Fraction of L2/3 PNs responding to long (200 ms, 30 mA) HP stimulation at onset only (ON; n = 2) and at both onset and offset (ON + OFF; n = 12). (B) Example traces of ON + OFF responses over 40 trials (gray) superimposed with the average response (black) for an individual L2/3 PN. (C) Grand average of ON + OFF responses (8 cells, 320 trials). (D) Distribution of onset latency, peak latency, half-amplitude duration, and amplitude (from left to right) of ON+OFF responses. Box plots shows the median, interquartile, range, and individual values. Statistical significance was calculated by paired Student’s t test. * P < 0.05.

## DISCUSSION

We describe the intrinsic properties, up- and down-states, spontaneous firing activity, and sensory stimulus evoked responses of layer 2/3 pyramidal neurons of the hind paw-related primary somatosensory cortex of anesthetized mice. We report a low spontaneous firing rate and a sparse sensory stimulus evoked firing code. Furthermore, approximately 50% of the recorded neurons did not respond to contralateral HP stimulus, and displayed a lower intrinsic excitability when compared with the responding neuronal population. Responses were characterized by long latencies following tactile stimulus, and, for stance-like stimulation, by the presence of both on- and off-responses.

Tactile sensory processing has been extensively described for the S1-BC. However, limited attention has been afforded to S1-HP responses. One striking feature of our findings for L2/3 pyramidal of the S1-HP region was the long latency of their stimulus-evoked responses compared with S1-BC (Crochet and Petersen, 2006; Crochet et al., 2011; Gambino et al., 2014; Hubatz et al., 2020; Jouhanneau et al., 2014) and FP stimulation in mice (Zhao et al., 2016), and may reflect the functional role of HP information for the animal. Whisker stimulation evokes a fast depolarizing response with ~9 ms latency in L2/3 pyramidal neurons (Brecht et al., 2003; de Kock et al., 2007; Higley and Contreras, 2005; Jouhanneau et al., 2014; Petersen et al., 2003), in coherence with the role of whiskers for sensory processing with high temporal precision. In contrast, tactile stimulation of the hind paw (~23 ms for HP stimulation as shown here) resulted in responses with longer latencies, likely reflecting the hind-paws’ main functional role for locomotion. It is possible that onset latencies are different in awake behaving mice as shown for L2/3 PNs of S1-FP (Milenkovic et al., 2014; Zhao et al., 2016). Our results, as well as those derived from forepaw tactile stimulation experiments (Milenkovic et al., 2014) suggest that paw-related sensory information processing operates on a different temporal scale when compared to whisker-related information. In contrast to the latencies, the response amplitude of S1-HP and S1-BC neurons was similar. HP stimulus evoked EPSPs of L2/3 pyramids of S1-HP had an average amplitude of ~9 mV (*Figure 2C*), which is comparable to the amplitude measured for L2/3 pyramids of the forepaw, whisker-related, and oral area of S1 in rodents (Brecht et al., 2003; Clemens et al., 2018; Crochet et al., 2011; Gambino et al., 2014; Higley and Contreras, 2005; Milenkovic et al., 2014; Zhao et al., 2016). Similarly, the response duration measured in our experiments for S1-HP was similar to that for L2/3 PNs of S1-BC and the oral S1 area (Clemens et al., 2018; Gambino et al., 2014).

For neurons of the supragranular layers of primary sensory cortices (somatosensory, visual, auditory) in rodents, many studies have described sparse firing activity — both spontaneously and following sensory stimulus (Brecht et al., 2003; Crochet et al., 2011; de Kock and Sakmann, 2009; Kerr et al., 2007; Kerr et al., 2005; Poulet and Petersen, 2008; Zhao et al., 2016). This sparse activity was substantiated by our study of the S1-HP region. Indeed, we found that the spontaneous firing activity was ~0.16 Hz for active cells, and that approximately 50% of the L2/3 PNs remained silent during the recordings. These values are similar to those reported for L2/3 PNs of primary somatosensory cortices (Barth and Poulet, 2012; Jouhanneau et al., 2014; Palmer et al., 2014), except for S1-FP L2/3 PNs that have higher spontaneous AP firing rates (Zhao et al., 2016). Similarly, hind paw stimulation elicited AP firing in only ~10% of L2/3 PNs that respond to HP stimulus, and these APs occurred in less than 50% of the trials with a maximum of one AP per trial. It should be noted, however, that the level of spontaneous and evoked firing activity likely depends on the brain state and the animal’s behavior (passive or active) (de Kock and Sakmann, 2008, 2009; Greenberg et al., 2008; Zhao et al., 2016).

The duration and frequency of up- and down-states in S1-HP L2/3 PNs described in our study are comparable to those described in the literature for rodent sensory cortices (Chance et al., 2002; Haider et al., 2007; Hasenstaub et al., 2007; Luczak et al., 2009; Palmer et al., 2014; Petersen et al., 2003; Zhao et al., 2016). During up-states S1-HP responses to brief tactile stimulation were reduced, similar to what has been described for S1 responses to whisker deflection as well as thalamic stimulations in rodents (Higley and Contreras, 2005; Petersen et al., 2003; Sachdev et al., 2004; Watson et al., 2008). Interestingly, sensory stimulus evoked sub-threshold responses are enhanced in the cat visual cortex during up-states (Haider et al., 2007). Whether these opposing findings are modality-, area, or species-specific features requires further investigation.

Our finding that the tactile stimulus used in our study triggered a response in only half of the L2/3 PNs population within S1-HP might support the idea that L2/3 contains sub-populations of pyramidal neurons with different functional roles (Chen et al., 2013; Peron et al., 2015; Yamashita et al., 2013). For instance, certain S1-HP sub-populations might be activated by other forms of vibrotactile stimuli, by temperature or painful stimuli (see (Milenkovic et al., 2014; Paricio-Montesinos et al., 2020). Different functional roles of these L2/3 PNs sub-populations are likely correlated with distinct brain-wide connectivity maps and intrinsic excitability properties (Chen et al., 2013; Yamashita et al., 2013). In agreement with these findings, we found that neurons that responded to hind-paw stimulation displayed an increased intrinsic excitability when compared to those that did not respond.

We demonstrated that the majority of L2/3 PNs responded to a prolonged paw stimulus with an on- and off component, reflecting the activation of the paw during the stance phase of locomotion. The off response could be a consequence of the activation of fast adaptive receptors in the glabrous skin of the HP when the stimulation ceases (Roudaut et al., 2012). Pacinian and Meissner corpuscles are both rapidly adapting mechanoreceptors that could underlie the off response (Roudaut et al., 2012; Zimmerman et al., 2014).

Apart from locomotory function, mouse utilizes its paws to explore, detect and discriminate various objects and sensory stimuli, which will allow them to perceive the environment and behave accordingly (Hirasawa et al., 2016; Morandell and Huber, 2017; Prsa et al., 2019; Zimmerman et al., 2014). These different functional roles are regulated by various receptor types present in the periphery and their associated neurons carrying the information to the brain. The neurons’ responses vary according to the way a peripheral receptor perceives a sensory stimulus and at what stimulus intensity they are being activated. For instance, rapidly adapting mechanoreceptors (RAMs) of the hind paws are less sensitive and less densely expressed compared to the RAMs of the forepaws, and are innervated with fewer axons. In contrast, slowly adapting mechanoreceptors (SAMs), which respond to low intensity threshold stimuli are comparable between fore- and hind paws (Walcher et al., 2018). The rodent’s paws also contain the most sensitive hair follicles with larger receptive fields, nociceptors for the noxious stimuli, and warm and cold receptors. The uniqueness in their tuning properties, adaptation rates and conduction velocity allow the paw system to respond to multiple sensory modalities. However, the afferent systems and central pathways to which these different receptors are connected are not completely understood (Abraira and Ginty, 2013; Arcourt et al., 2017; Milenkovic et al., 2014; Paricio-Montesinos et al., 2020; Walcher et al., 2018; Zimmerman et al., 2014).

SAMs that encode touch and are densely expressed in the mouse forepaw are also highly innervated in the human finger pad coding for stimulus orientation and discrimination in humans (Hsiao et al., 2002; Walcher et al., 2018). Notably, distinct receptors in the fingertips and their complementary neural circuits are involved in different tasks like tactile roughness discrimination or spatial resolution acuity (Libouton et al., 2010). Similarities in arrangements of touch receptors across the rodent paws and existence of identical receptors and neuronal pathways in primates and human limbs makes this system highly translational compared to the well-established whisker tactile system in rodents (Abraira and Ginty, 2013; Johnson, 2001; Leem et al., 1993; Walcher et al., 2018). It is crucial to define a task which is applicable both to a laboratory animal model and human beings. Our work describes relevant characteristic features of a sensory response to paw stimulation in L2/3 pyramidal neurons of the S1-HP that are shared with other tactile modalities as well as unique for the paw related information. This work opens new avenues for comparing tactile sensory responses in L2/3 PNs of WT mice with those of mouse models of sensory processing defects, or for exploring developmental changes in sensory processing.

## ADDITIONAL INFORMATION

### Author contribution

GB and AAB carried out electrophysiological recordings, data analysis and staining of neurons. KLC carried out staining of neurons. AF conceived the project and designed experiments. All authors wrote the manuscript and have approved its final version. All persons designated as authors qualify for authorship, and all those who qualify for authorship are listed.

## Acknowledgements

We would like to thank Dr. M. Brecht for help with setting up the *in vivo* patch-clamp techniques in our laboratory. We are also grateful to Dr. M. Ginger for fruitful discussions and feedback on the manuscript. The images were acquired using equipment of the Bordeaux Imaging Center.

