## Supplementary material for "Sensory stimulus evoked responses in layer 2/3 pyramidal neurons of the hind paw-related mouse primary somatosensory cortex": Supple Figure 1

**Supplementary figure**

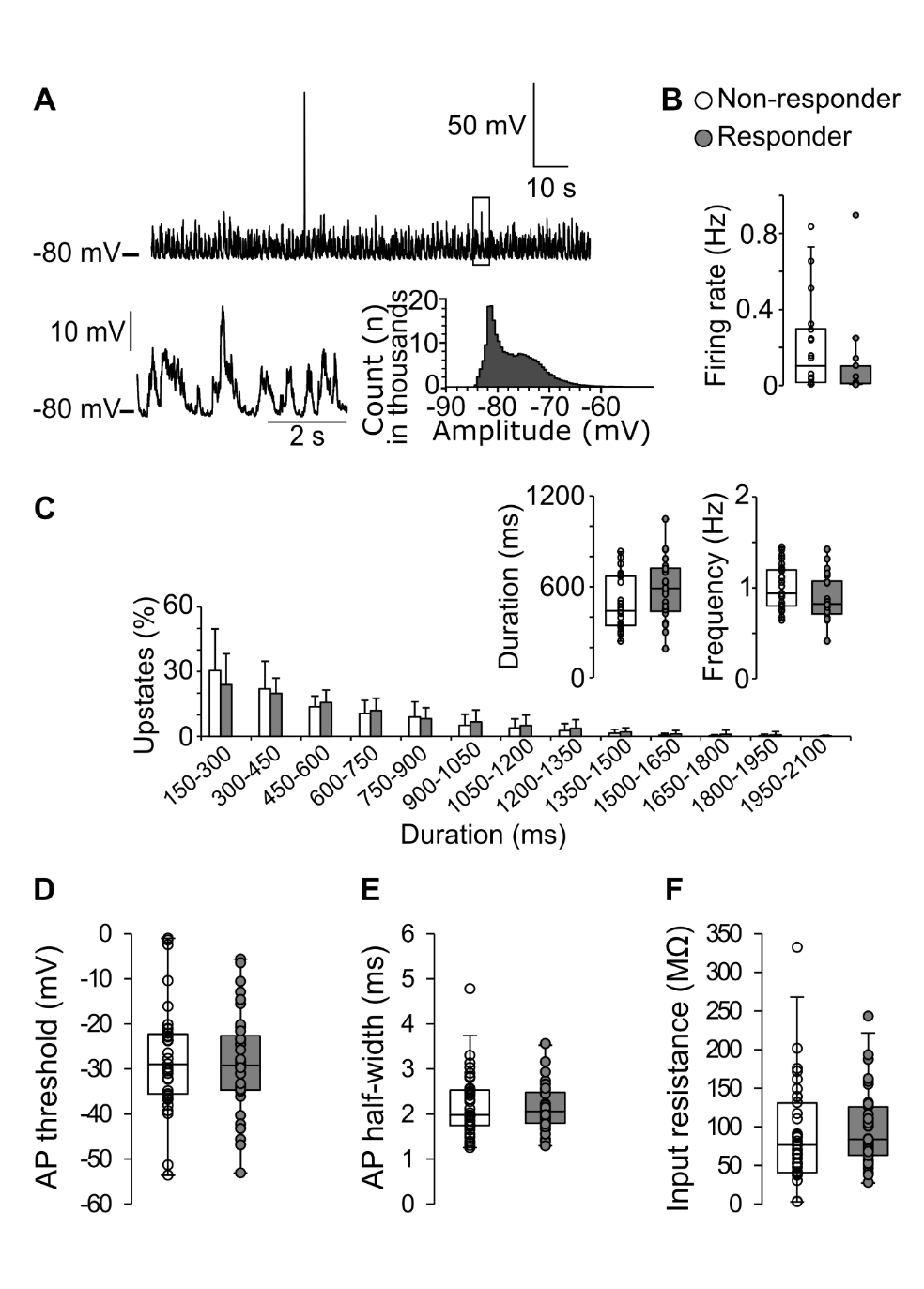

***Supplementary figure 1. Spontaneous firing activity and slow wave oscillations in L2/3-pyramidal neurons.*** *(A) Example of a 2-min continuous recording of spontaneous firing activity and up- and down-states in a Responder cell (top). Zoom-in of a 5-sec period (see box) of the example shown above (bottom left). Plot of membrane potential distribution (bottom right). (B) Spontaneous firing rate for actively firing Responders and Non-responders. (C) Population data for up-states. Plot of the fraction of up-state durations, as well as average duration and frequency (insets). (D) Average AP threshold, (E) AP half-width, and (F) input resistance for R- (n = 37) and NR-cells (n = 35). Box plots show the median, interquartile, range, and individual values. Statistical significance was calculated by two-way ANOVA with repetition or unpaired Student’s t test.*
